# The YAP-TEAD complex promotes senescent cell survival by lowering endoplasmic reticulum stress

**DOI:** 10.1101/2022.12.18.520921

**Authors:** Carlos Anerillas, Krystyna Mazan-Mamczarz, Allison B. Herman, Rachel Munk, Kwan-Wood Gabriel Lam, Miguel Calvo-Rubio, Amanda Garrido, Dimitrios Tsitsipatis, Jennifer L. Martindale, Gisela Altés, Martina Rossi, Yulan Piao, Jinshui Fan, Chang-Yi Cui, Supriyo De, Kotb Abdelmohsen, Rafael de Cabo, Myriam Gorospe

**Affiliations:** Laboratory of Genetics and Genomics, National Institute on Aging Intramural Research Program, National Institutes of Health, Baltimore, Maryland, USA; Translational Gerontology Branch, National Institute on Aging Intramural Research Program, National Institutes of Health, Baltimore, Maryland, USA

## Abstract

Sublethal cell damage can trigger a complex adaptive program known as senescence, characterized by growth arrest, resistance to apoptosis, and a senescence-associated secretory phenotype (SASP). As senescent cells accumulating in aging organs are linked to many age-associated diseases, senotherapeutic strategies are actively sought to eliminate them. Here, a whole-genome CRISPR knockout screen revealed that proteins in the YAP-TEAD pathway influenced senescent cell viability. Accordingly, treating senescent cells with a drug that inhibited this pathway, Verteporfin (VPF), selectively triggered apoptotic cell death and derepressed DDIT4, in turn inhibiting mTOR. Reducing mTOR function in senescent cells diminished endoplasmic reticulum (ER) biogenesis, causing ER stress and apoptosis due to high demands on ER function by the SASP. Importantly, VPF treatment decreased senescent cell numbers in the organs of old mice and mice exhibiting doxorubicin-induced senescence. We present a novel senolytic strategy that eliminates senescent cells by hindering ER activity required for SASP production.

## INTRODUCTION

Cellular senescence is a dynamic state of cells responding to sublethal damage, characterized by persistent growth arrest and a secretory activity known as the senescence-associated secretory phenotype (SASP) (Gorgoulis et al., 2019). Other prominent traits of senescent cells include an altered metabolic profile, resistance to apoptosis, and enduring damage to DNA and other macromolecules. Although cellular senescence can be beneficial during developmental processes like embryonic morphogenesis, tissue repair, and cancer prevention, the aberrant accumulation of senescent cells is detrimental and leads to tissue decline and disease, as seen during the aging process (Di Micco et al., 2021). Recently characterized compounds known as senolytics aim to reduce the accumulation of senescent cells within tissues (Gasek et al., 2021). Most of the known senolytics exploit the fact that senescent cells have endured severe damage but remain alive due to the implementation of robust anti-apoptotic programs. Accordingly, many senolytics function by disrupting this balance in senescent cells, either by interfering with pro-survival factors or by promoting the actions from pro-apoptotic factors (Baar et al., 2017; Kirkland and Tchkonia, 2020).

To identify new strategies for removing senescent cells, we performed a whole-genome CRISPR screen to identify genes that function to preserve the viability of senescent cells. We report the discovery of several genes encoding proteins in the Hippo pathway, which modulates the transcriptional activity of the YAP-TEAD complex, as being essential for maintaining senescent cell survival. Moreover, we found that treatment with a drug named verteporfin (VPF), that inhibits the interaction between YAP and TEAD and thus prevents the transcriptional function of this complex, caused senolysis in several cell culture models of senescence. Treatment of senescent cells with VPF triggered endoplasmic reticulum (ER) stress and subsequent apoptosis. Detailed characterization of VPF actions revealed that, in senescent cells, YAP-TEAD repressed the transcription of DDIT4, an inhibitor of mTOR (Lipina and Hundal, 2016); accordingly, treatment with VPF reduced mTOR function and lowered the biosynthesis of the ER wall. We propose a model whereby interfering with the YAP-TEAD—DDIT4—mTOR pathway renders senescent cells more susceptible to senolysis due to their unique requirement for robust protein synthesis to sustain the SASP. In support of this model, ablating the SASP by silencing RELA rescued senescent cells from the apoptotic impact of inhibiting YAP-TEAD or mTOR. Finally, VPF treatment reduced the burden of senescent cells in live organisms, as seen in naturally aging mice and mice exhibiting doxorubicin-induced senescence. Considering these findings, we propose that metabolic targeting of the secretory machinery can sensitize senescent cells to achieve senolysis.

## RESULTS

### CRISPR screen uncovers a key role for the YAP-TEAD pathway in senescent cell viability

A CRISPR screen was devised to identify pathways that sensitize senescent cells to senolysis. Briefly, we triggered etoposide-induced senescence (ETIS) in WI-38 human diploid fibroblasts by treatment with 50 μM etoposide following the scheme indicated (**Figs. 1A, B**). After confirming the presence of senescent cells by assessing senescence-associated β-Galactosidase (SA-β-Gal) activity and BrdU incorporation (**Figs. 1C-E**), we transduced ETIS WI-38 cells with a human, whole-genome CRISPR knockout library (Brunello) (Doench et al., 2016) at a multiplicity of infection (MOI) of 0.44 after optimization (**Fig. S1A**). Seventy-two hours later, we collected a reference sample (t=0), and incubated cells for an additional 14 days before analysis to ensure identification of gRNAs with slow kinetics. To identify genes relevant for viability, we focused on gRNAs showing decreased representation after 14 days (**Fig. 1F; Supplementary Tables 1, 2**). Enrichr analysis revealed that the Hippo/MST1 pathway, including the YAP-TEAD complex, were present at the top of multiple ranks such as Gene Ontology (GO) Biological Process, Reactome, or Cellular Component (**Figs. 1G, H**). We validated these results by individually silencing those genes in the Hippo YAP-TEAD pathway that were reduced in the CRISPR screen (*YAP1, TEAD2, MOB1A, MAP4K1*, and *TAOK2*). Among them, silencing *TEAD2, MOB1A*, and *MAP4K1* mRNAs reduced the number of viable senescent cells (**Figs. S1B, C**). Western blot analysis revealed that the levels of Hippo components were unchanged (**Fig. S1D**), but the activity of the YAP-TEAD complex increased in senescent cells, as a heterologous luciferase reporter driven by a TEAD-responsive promoter and the levels of YAP-TEAD transcriptional targets *ANKRD1* and *TGFB2* mRNAs (Dupont et al., 2011; Lee et al., 2016; Srikantan et al., 2011) increased in senescent cells (**Figs. 1I, J**). Moreover, silencing YAP1 or TEAD2 reduced the levels of *ANKRD1* and *TGFB2* mRNAs back to control levels, while the levels of *p16* mRNA, encoding a senescence marker (Munoz-Espin and Serrano, 2014), remained elevated (**Fig. 1J**).

**Figure 1.**
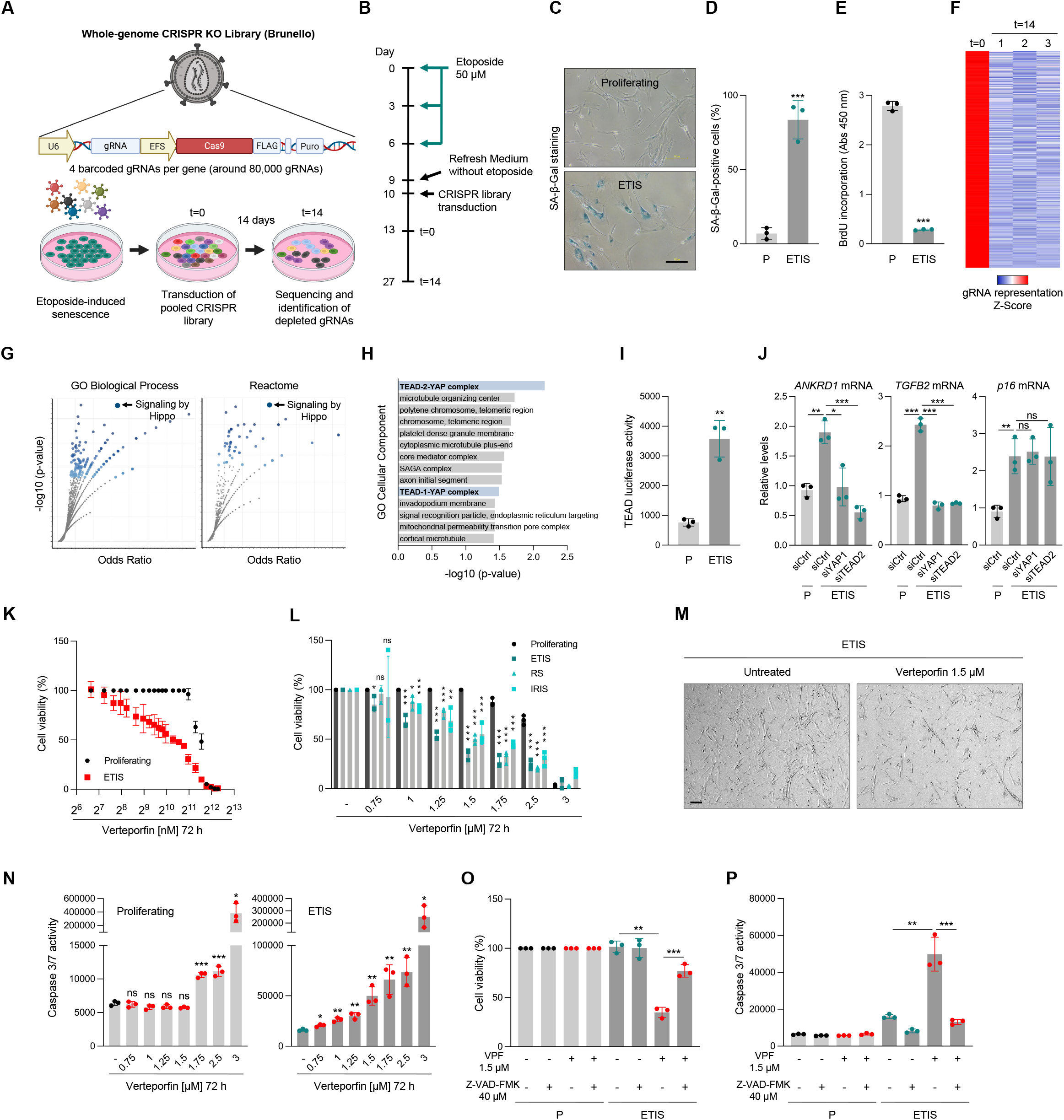
A CRISPR screen identifies YAP-TEAD pathway as essential for senescent cell viability. **(A)** Schematic of the CRISPR screen performed on etoposide-induced senescent (ETIS) WI-38 fibroblasts, including the lentiviral construct transduced and the different steps taken. **(B)** Timeline of the treatments and procedures in (A). **(C,D)** Representative images of senescence-associated-β-galactosidase staining (C) and quantification (D) in proliferating (P) and ETIS WI-38 cells. Scale bar, 100 μm. **(E)** BrdU assay performed in the conditions described in (C). **(F)** Heatmap of the Z-Scores of the gRNAs that were significantly reduced when comparing t=14 to t=0 (further information in Supplementary Table 1). **(G)** Analysis of the gRNAs depleted from the t=14 experimental groups (STAR Methods) with Enrichr (further information in Supplementary Table 2). The dot plots show the combined score (y and x axes) of “Signaling by Hippo” categories from GO and Reactome databases. For each category displayed (dots present in the plot) the *y* axis represents their -log10(p-value) while the *x* axis represents the odds ratio. **(H)** Bar plot showing the combined scores of GO database “Cellular Component” obtained by Enrichr analysis of the conditions described in (G). **(I)** Luciferase activity of a TEAD reporter construct analyzed for the indicated experimental groups. **(J)** Measurement by RT-qPCR analysis of the indicated mRNAs (normalized to *ACTB* mRNA) in ETIS WI-38 transfected with siCtrl, siYAP1, or siTEAD2; 24 h later, cells were treated with etoposide (50 μM) for 8 days. Proliferating cells transfected with siCtrl were included as a control. **(K)** Cell viability assessment by direct cell counting of the indicated Verteporfin doses in either proliferating or ETIS WI-38 fibroblasts. **(L)** Cell viability analysis by direct cell counting of the indicated Verteporfin (VPF) doses in WI-38 fibroblasts that were proliferating or rendered senescent by ETIS, replicative senescence (RS), or IR-induced senescence (IRIS). **(M)** Representative micrographs of senescent (ETIS) WI-38 cells that were either untreated or treated with 1.5 μM Verteporfin for 72 h. Scale bar, 100 μm. **(N)** Caspase 3/7 activity measurement in either proliferating or senescent (ETIS) WI-38 cells with the indicated doses of Verteporfin. **(O, P)** Cell viability evaluation by direct cell counting (O) and Caspase 3/7 activity measurement (P) in Proliferating and ETIS WI-38 cells treated with Verteporfin (VPF), in which apoptosis was rescued by simultaneous treatment with Z-VAD-FMK where indicated. Graphs in (D, E, I-L, N-P) display the means and each individual value as a dot ±SD of at least n=3 independent replicates; significance (*P< 0.05, **P< 0.01, ***P< 0.001) was determined using two-tailed Student’s *t*-test. Unless indicated, statistical tests were performed relative to untreated or proliferating controls. See also Fig. S1.

We then tested in senescent and proliferating WI-38 cells the impact of the drug verteporfin (VPF), which functions by disrupting the interaction between YAP and TEAD (Liu-Chittenden et al., 2012). As shown in **Fig. 1K**, after incubation with VPF at varying concentrations for 72 h, senescent cells were significantly more sensitive than proliferating cells to VPF. We extended this analysis to other models of senescence, including WI-38 fibroblasts undergoing replicative senescence (RS) and rendered senescent following exposure to ionizing radiation (15 Gy followed by 10 days in culture; IRIS); senescence was confirmed by assessing SA-β-Gal activity and BrdU incorporation (**Figs. S1E, F**). In these senescence models, VPF treatment selectively reduced the viability of senescent cells (**Figs. 1L, M**). As in previous studies evaluating senescent cell viability by monitoring caspase activity (Anerillas et al., 2022a; Munk et al., 2021), caspase 3/7 activity increased specifically in senescent cells treated with up to 1.5 μM VPF (**Fig. 1N, Fig. S1G**), while higher VPF doses increased caspase 3/7 activity also in proliferating cells. To confirm that VPF triggered cell death in senescent cells through apoptosis, cell death was rescued by blocking caspase 3/7 activity with the pan-caspase inhibitor Z-VAD-FMK (**Figs. 1O,P**). Similarly, VPF triggered senolysis in other human cell types [diploid fibroblasts BJ and IMR-90, small airway epithelial cells (HSAEC), and umbilical vein endothelial cells (HUVEC)] rendered senescent by treatment with etoposide (**Figs. S1H, I**). Together, these data suggest that senescent cells can be selectively eliminated by inhibiting YAP-TEAD function.

### Inhibiting YAP-TEAD triggers apoptosis by derepressing DDIT4, causing ER stress

To investigate how inhibition of YAP-TEAD by VPF triggered apoptosis in senescent cells, we studied the effects of VPF at 48 h, when signs of cell death began, to understand the signaling changes leading to apoptosis. First, we confirmed that VPF reduced the interaction between YAP and TEAD and hence YAP-TEAD function in this paradigm. As shown in **Fig. 2A**, VPF treatment reduced YAP-TEAD luciferase activity, decreased the levels of transcriptional YAP-TEAD targets *ANKRD1, TGFB2, WNT5B, LATS2, NF2, ADAMTS1*, and *AMOTL1* mRNAs (**Fig. S2A**) (Stein et al., 2015; Zanconato et al., 2015), and hindered the interaction of YAP and TEAD as observed by coimmunoprecipitation (co-IP) analysis (TEAD IP followed by western blot analysis to detect YAP levels in the IP; **Fig. S2B**).

**Figure 2.**
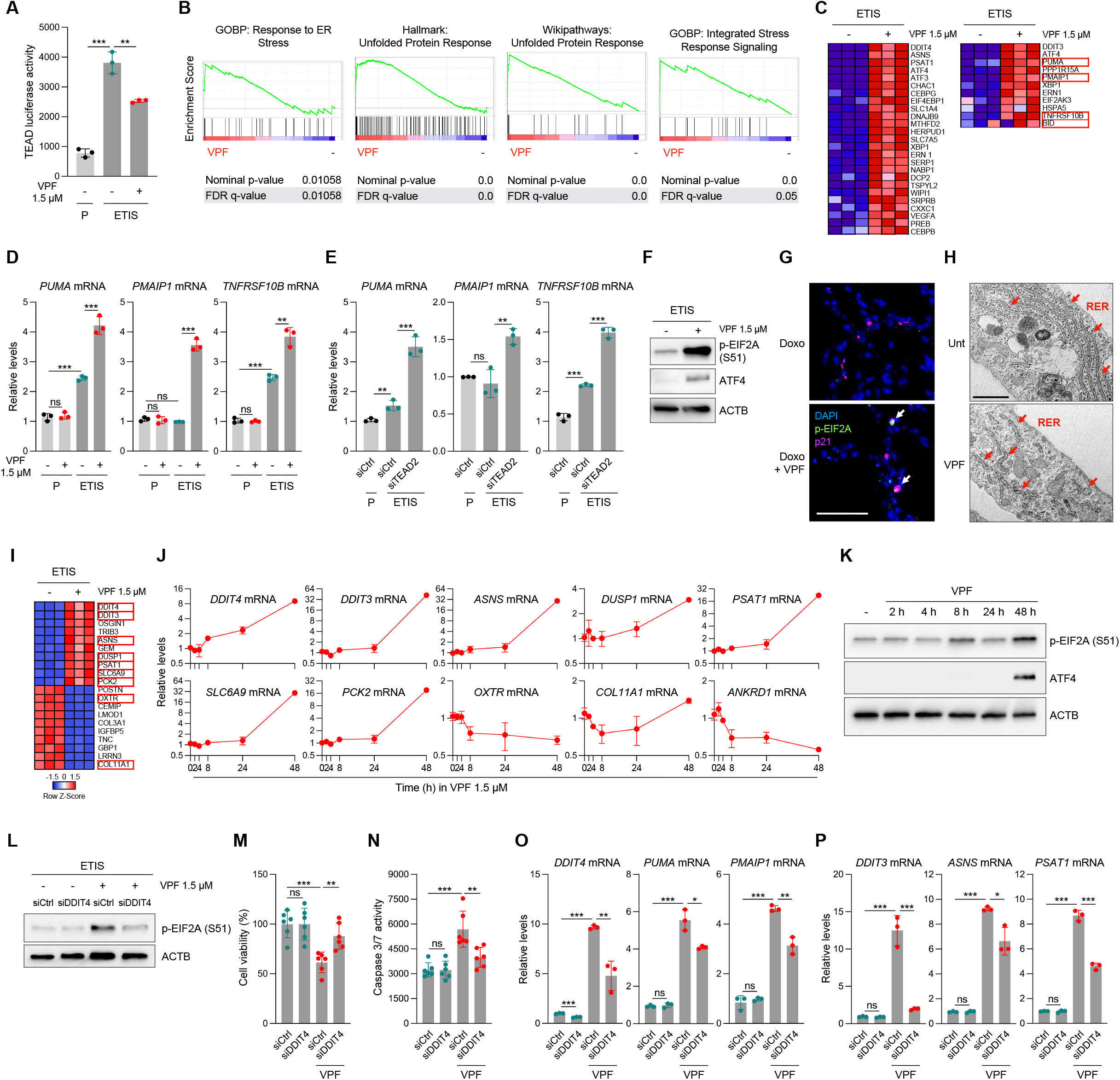
YAP-TEAD inhibition triggers apoptosis by derepressing DDIT4, causing ER stress. **(A)** Luciferase activity from the TEAD promoter as measured in ETIS WI-38 cells that were either left untreated or treated with 1.5 μM VPF for 48 h starting at day 6 after etoposide treatment; proliferating (P) cells were included as a control population. **(B)** GSEA analysis of RNA-seq performed in ETIS WI-38 cells treated with VPF relative to control untreated cells as in (A). Plots represent the transcriptomic association of the VPF-treated ETIS WI-38 cells with the indicated gene sets, as compared to untreated ETIS WI-38; GOBP: GO Biological Process. **(C)** Heatmap displaying the levels of each of the transcripts included in the GSEA gene sets shown (Hallmark: Unfolded Protein Response, left heatmap; WikiPathways: Unfolded Protein Response, right heatmap) for the conditions described in (B). mRNAs encoding pro-apoptotic proteins are highlighted by red boxes. **(D)** Quantification by RT-qPCR analysis of the indicated transcripts in P and ETIS WI-38 cells that were either untreated or treated with VPF for 48 h as in (A). **(E)** Quantification by RT-qPCR analysis of the indicated transcripts in ETIS WI-38 cells transfected with siCtrl or siTEAD2 one day before senescence induction. Proliferating WI-38 cells transfected with siCtrl were used as a control. **(F)** Representative western blot analysis of the levels of phosphorylated EIF2A (S51), ATF4, and loading control ACTB in WI-38 cells treated as described in (B). **(G)** Representative images of immunofluorescence performed in lungs from mice that had been injected with doxorubicin 14 days earlier and were treated daily with either DMSO or VPF (50 mg/kg) from day 10 onward (see also Fig. S4A). Cells were stained to visualize nuclei (DAPI, blue), phosphorylated-EIF2A (S51; green) and p21 (purple); double stained cells are indicated with a white arrow. Scale bar, 50 μm. **(H)** Representative images of transmission electron microscopy (TEM) of ETIS WI-38 cells treated with VPF as in (A), along with untreated controls (Unt). Red arrows point to the rough endoplasmic reticulum (RER) observed. Scale bar, 1 μm. **(I)** Heatmap displaying the row Z-Score of each of the transcripts indicated (top 10 increased and decreased in VPF-treated ETIS WI-38 cells compared to untreated ETIS WI-38 controls) between conditions. Red boxes indicate that the mRNA is a reported direct target of the YAP-TEAD complex and is also part of the ER stress response. **(J)** RT-qPCR analysis of the indicated YAP-TEAD target mRNAs at the indicated times after treatment of senescent (ETIS) WI-38 cells with 1.5 μM VPF. **(K)** Western blot analysis of the levels of phosphorylated EIF2A (S51), ATF4, and loading control ACTB in cells treated as described in (J). **(L)** Western blot analysis of phosphorylated EIF2A (S51) and loading control ACTB levels in ETIS WI-38 cells transfected with the indicated siRNAs, and then either untreated or treated with 1.5 μM VPF for 48 h. **(M, N)** Assessment of cell viability by direct cell counting (M) and Caspase 3/7 activity (N) for the experimental groups described in (L). **(O, P)** RT-qPCR analysis of the indicated mRNAs in the experimental groups described in (L). Graphs in (A, D, E, J, M-P) show the means and specify each of the individual values as a dot ±SD of at least n=3 independent replicates; significance (*P< 0.05, **P< 0.01, ***P< 0.001) was calculated by performing two-tailed Student’s *t*-test. See also Fig. S2.

We then performed RNA-sequencing (RNA-seq) analysis of untreated and VPF-treated senescent cells (ETIS) and employed Gene Set Enrichment Analysis (GSEA) to identify differences in the two transcriptomes (GSE221104, token wpexgsycrbqhdcn). As shown in **Fig. 2B**, most of the top-ranked GSEA gene sets associated with VPF treatment were related to the endoplasmic reticulum (ER) stress response, including the Unfolded Protein Response (UPR) and the Integrated Stress Response (ISR), both of which converge on the central effector eIF2α (EIF2A) (Pakos-Zebrucka et al., 2016). In line with recent reports (Chan et al., 2022), we found that the Epithelial-Mesenchymal Transition (EMT) signature in senescent cells was reduced by VPF treatment (**Fig. S2C**). We focused on ER stress given its predominant representation in pathway analysis and its strong link to apoptosis (Tabas and Ron, 2011), as supported by the increased levels of mRNAs encoding pro-apoptotic proteins like PUMA, PMAIP1, TNFRSF10B, and BID (Anerillas et al., 2022b; Baar et al., 2017) (**Fig. 2C,** red rectangles). These observations were validated by RT-qPCR analysis in both VPF-treated and TEAD2-silenced senescent WI-38 cells (**Figs. 2D, E**).

To further characterize the mediators of ER stress caused by VPF in senescent cells, we focused on the ER stress response factors EIF2A and ATF4, as these proteins are shared effectors of the ISR and the UPR (Tabas and Ron, 2011). As shown, the levels of phosphorylated (p-) EIF2A and ATF4 increased in VPF-treated senescent cells (**Fig. 2F**), but the levels of XBP1s and ATF6 (**Fig. S2D**), which are key effectors of alternative ER stress responses (Hetz and Papa, 2018), did not. GSEA analysis also pointed to PERK (EIF2AK3), an upstream activator of EIF2A previously related to senescence (Zhang et al., 2022), as a prominent contributor to the transcriptomic signature displayed by VPF-treated senescent cells (**Fig. S2E**). In fact, PERK silencing reduced both EIF2A phosphorylation and cell death (**Fig. S2F-H**) caused by VPF treatment in senescent cells. Moreover, VPF treatment caused ER stress in lungs from mice in which senescence was triggered by treatment with one dose of doxorubicin (10 mg/kg) and analyzed 14 days later (**Fig. 2G**) (Anerillas et al., 2022b; Demaria et al., 2017). Finally, analysis of VPF-treated senescent (ETIS) WI-38 fibroblasts by transmission electron microscopy (TEM) showed that the rough endoplasmic reticulum (RER) of these cells appeared more swollen and less organized, two signs of ER stress commonly assessed using this technique (Montalbano et al., 2013; Oslowski and Urano, 2011), than that of untreated senescent cells (**Fig. 2H**). Together, these data suggest that VPF treatment causes apoptosis by signaling through the ER stress response factors PERK, EIF2A, and ATF4.

Since the YAP-TEAD complex can both enhance and repress transcription (Kim et al., 2015; Stein et al., 2015; Zanconato et al., 2015), we examined the top 10 mRNAs that increased or decreased with VPF and focused on known direct targets of YAP-TEAD, such as *DDIT4, DDIT3, ASNS, DUSP1, PSAT1, SLC6A9, PCK2, OXTR*, and *COL11A1* mRNAs (**Fig. 2I**). We then assessed the kinetics of expression of each mRNA, including *ANKRD1* mRNA as a control readout of YAP-TEAD function, following addition of VPF (**Figs. 1J** and **S2A**). We observed that only *DDIT4, OXTR*, and *COL11A1* mRNAs, along with the control, *ANKRD1* mRNA, changed significantly by 8 h (**Fig. 2J**). Interestingly, *DDIT4* mRNA, which is transcriptionally repressed by YAP-TEAD (Kim et al., 2015), is part of the ER stress response transcriptomic program (Pakos-Zebrucka et al., 2016) (**Fig. 2C, *left***), and appeared to increase before other ER stress-related transcripts (*DDIT3, ASNS*, and *PSAT1* mRNAs). Since the levels of ATF4 (and p-EIF2A) decline by 48 h of VPF treatment (**Fig. 2K**), when *DDIT3, ASNS*, and *PSAT1* mRNAs are strongly increased (**Fig. 2J**), the derepression of *DDIT4* mRNA might represent an earlier event that later leads to a full ER stress response. Chromatin IP (ChIP) analysis indicated that VPF treatment reduced the association of YAP to a binding region in the *DDIT4* gene promoter (Kim et al., 2015) (**Fig. S2K**). These findings support the hypothesis that *DDIT4* mRNA derepression is a direct consequence of YAP-TEAD inhibition rather than an indirect result of the ER stress response.

As p53 controls *DDIT4* mRNA levels transcriptionally (Tirado-Hurtado et al., 2018), we hypothesized that DDIT4 derepression could be mitigated by silencing p53 in VPF-treated senescent cells. However, as shown in **Fig. S2L**, *DDIT4* mRNA levels were lower in senescent cells compared to proliferating cells, even though p53 activity is elevated in senescence, and p53 silencing reduced *DDIT4* mRNA levels in VPF-treated senescent cells. These data suggest that YAP-TEAD function could counteract some pro-apoptotic features of p53 in senescent cells, as observed for *DDIT4* mRNA. Finally, to confirm that DDIT4 contributes to ER stress and subsequent senolysis in VPF-treated senescent cells, we silenced DDIT4 before treatment with VPF. Reducing DDIT4 levels decreased both p-EIF2A levels and apoptotic cell death caused by VPF (**Figs. 2L-N**). Accordingly, the rise in mRNAs related to the ER stress response and apoptosis by VPF was diminished after silencing DDIT4 (**Figs. 2O,P**). In sum, ER stress and senolysis caused by VPF appear to be mediated by DDIT4 derepression.

### Inhibition of mTOR-dependent ER biogenesis by DDIT4 induces ER stress and senolysis

Given that DDIT4 inhibits mTOR (Lipina and Hundal, 2016), we analyzed if VPF treatment affected signaling through PI3K-AKT-mTOR in senescent cells. In agreement with the known function of DDIT4 as an inhibitor of RHEB (Foltyn et al., 2019; Lipina and Hundal, 2016), we found that mTOR and S6K phosphorylation were reduced by VPF treatment, leaving AKT unaffected (**Fig. 3A**). Moreover, VPF treatment inhibited mTOR after only 8 h (**Fig. 3B**), coinciding with the derepression of *DDIT4* mRNA (**Fig. 2J**) and before the full implementation of ER stress-related increase in EIF2A phosphorylation and ATF4 levels (**Fig. 2K**). Silencing DDIT4 significantly mitigated the inhibition of S6K phosphorylation caused by VPF (**Fig. 3C**), suggesting that DDIT4 might also act as an inhibitor of mTOR in our system. In sum, mTOR inhibition by DDIT4 preceded ER stress in senescent cells treated with VPF.

**Figure 3.**
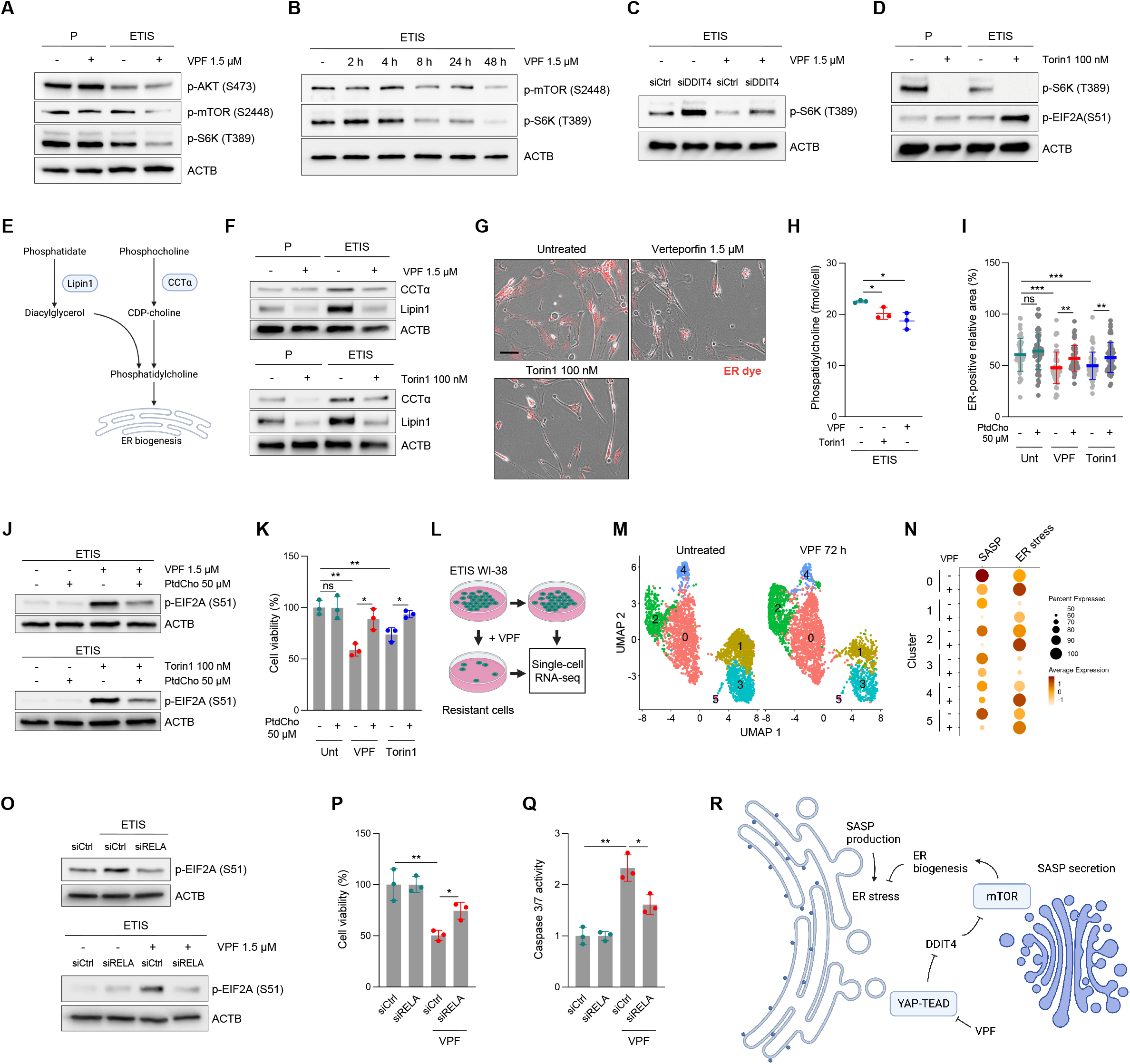
Inhibition of mTOR-dependent ER biogenesis by DDIT4 induces ER stress and senolysis. **(A)** Representative Western blot analysis of the levels of phosphorylated AKT (S473), phosphorylated mTOR (S2448), phosphorylated p70 S6K (T389), and loading control ACTB in proliferating (P) or etoposide-induced senescent (ETIS) WI-38 cells, either untreated or treated with 1.5 μM VPF for 48 h, starting at day 6 of etoposide treatment. **(B)** Western blot analysis of the levels of phosphorylated mTOR (S2448), phosphorylated p70 S6K (T389), and loading control ACTB at the indicated time points after treatment with 1.5 μM VPF. **(C)** Western blot analysis of the levels of phosphorylated p70 S6K (T389) and loading control ACTB in WI-38 cells transfected with siCtrl or siDDIT4, rendered senescent with etoposide for 8 days, and either left untreated or treated with 1.5 μM VPF for an additional 48 h. **(D)** Western blot analysis of the levels of phosphorylated p70 S6K (T389), phosphorylated EIF2A (S51), and loading control ACTB in P and ETIS WI-38 cells that were either left untreated or treated with 100 nM Torin1 for 48 h, starting at day 6 of etoposide treatment. **(E)** Schematic depicting mTOR-regulated enzymes Lipin1 and CCTα, which regulate the biosynthesis of phosphatidylcholine and subsequent ER biogenesis. **(F)** Western blot analysis of the levels of CCTα, Lipin1, and loading control ACTB, after treatment of P or ETIS WI-38 cells, with 1.5 μM VPF (*top*) or 100 nM Torin1 (*bottom*) for 48 h, as in (A, D). **(G)** Representative micrographs depicting the relative ER area stained with an ER tracker (in red) in ETIS WI-38 cells that were either untreated, treated with 1.5 μM VPF, or treated with 100 nM Torin1 for 48 h. Scale bar, 100 μm. **(H)** Dot plot representation of the levels of Phosphatidylcholine (PtdCho) (fmol/cell) as measured in the conditions described in (G). **(I)** Dot plot representation of the calculated ER-positive areas per cell for the experimental groups described in (G). Each of these treatments included simultaneous supplementation with 50 μM PtdCho. **(J)** Western blot analysis of phosphorylated EIF2A (S51) and ACTB levels with the indicated treatments, all of which were performed for 48 h in ETIS WI-38 cells. **(K)** Cell viability assessment by direct cell counting for the same groups described in (I). **(L)** Schematic overview of the experiment to identify single-cell transcriptomic differences between ETIS WI-38 without further treatment or treated with VPF for 72 h to cause a significant decrease of those cells most sensitive to VPF. **(M)** UMAP of single-cell gene expression showing six subgroups in ETIS WI-38 cells (left) and ETIS WI-38 cells subjected to 1.5 μM VPF for 72 h (right). **(N)** Dot plot showing the association of clusters described in (M) with ER stress (GOBP: Response to ER stress), apoptosis (Hallmark: Apoptosis), and SASP (a custom gene set of 132 markers) transcripts. Dot size corresponds to proportion of cells expressing transcripts in a gene set, and dot color represents the scaled expression level. **(O)** Representative Western blot analysis of the levels of phosphorylated EIF2A (S51) and loading control ACTB in proliferating and ETIS WI-38 cells treated with the indicated combinations of siRNAs and 1.5 μM VPF for 48 h. siCtrl-transfected, proliferating controls were also included. **(P, Q)** Cell viability assessment by direct cell counting (P) and Caspase 3/7 activity measurement (Q) in ETIS WI-38 cells after treatment for 48 h with the indicated combinations of siRNAs and VPF. **(R)** Schematic representing the proposed model for the role of YAP-TEAD in supporting the viability of senescent cells. Briefly, YAP-TEAD activity represses DDIT4 expression, which enables the mTOR activity necessary for ER biogenesis required to cope with the underlying ER stress caused by SASP production, thus preventing apoptosis. This tight balance can be disrupted by inhibiting YAP-TEAD with VPF, in turn causing senolysis. Graphs in (H, I, K, P, Q) show the means and each individual value as a dot ±SD of at least n=3 independent replicates; significance (*P< 0.05, **P< 0.01, ***P< 0.001) was calculated using two-tailed Student’s t-test. See also Fig. S3.

These observations prompted us to test if mTOR inhibition could cause ER stress in senescent cells. First, we tested the mTOR inhibitor Torin1, previously used in senescent cells (Herranz et al., 2015; Laberge et al., 2015), but we employed the higher dose of 100 nM, which did not reduce the viability of proliferating cells but caused significant death in senescent cells (**Figs. S3A, B**), as reported (Carroll et al., 2017). As shown, Torin1 treatment caused significant phosphorylation of EIF2A in senescent cells, along with S6K dephosphorylation (**Fig. 3D**); as seen in VPF-treated cells, this was the only branch of ER stress response triggered by Torin1, as the levels of ATF6 and XBP1s were unchanged by mTOR inhibition (**Fig. S3C**).

We sought a deeper understanding of how mTOR inhibition caused ER stress in senescent cells. YAP and TEAD support the ER enlargement required in cells with high demand for protein production (Wu et al., 2015) and mTOR is required for ER biogenesis (Jacquemyn et al., 2017; Mossmann et al., 2018). Therefore, we wondered whether mTOR inhibition caused by VPF might trigger ER stress by reducing the size of the ER. mTOR controls ER biogenesis at least in part by increasing the expression of two key enzymes in the production of phosphatidylcholine (PtdCho), Lipin1 (LPIN1) and CCTα (PCYT1A) (Brandt et al., 2018; Peterson et al., 2011; Quinn et al., 2017) (**Fig. 3E**). Importantly, treatment with VPF or Torin1 decreased Lipin1 and CCTα levels in senescent cells (**Fig. 3F**). To measure the relative size of the ER for each condition, we used a commercial ER tracker (ER-Tracker, STAR Methods) to quantify the relative areas positive for the dye in multiple cells from the different treatments; as shown, treatment with VPF or Torin1 significantly reduced the red area, a surrogate measure of ER size (**Fig. 3G, Fig. S3E**). Each treatment also reduced the levels of PtdCho per cell (**Fig. 3H**), suggesting that these drugs might lower the production of PtdCho, necessary for ER biogenesis. In further support of this hypothesis, we observed that the reduction in ER triggered by treatment with VPF or Torin1 was partially rescued by PtdCho supplementation (**Fig. 3I**, **Fig. S3F**). Moreover, both EIF2A phosphorylation and cell death caused by VPF or Torin1 were mitigated by PtdCho supplementation (**Figs. 3J,K**). These observations support the notion that active YAP-TEAD preserves mTOR function, in turn enhancing ER biogenesis and avoiding ER stress and apoptosis.

mTOR function was proposed to support the SASP (Herranz et al., 2015; Laberge et al., 2015) and mTOR inhibition is widely accepted as a senomorphic intervention (Soto-Gamez and Demaria, 2017). Therefore, we tested whether mTOR inhibition by VPF or Torin1 could reduce the levels of SASP and the impact of the reduced SASP load on ER stress. Strikingly, neither treatment (VPF or Torin1) reduced the levels of SASP mRNAs (**Fig. S3G**) or SASP factors in conditioned media (**Fig. S3H**). These observations could be explained by the higher doses of Torin1 used in our study [four times higher doses than used previously (Herranz et al., 2015; Laberge et al., 2015)], or by the fact that mTOR inhibition was not reported to cause complete SASP depletion when tested in fully senescent cells (Herranz et al., 2015; Laberge et al., 2015), as we did here. In other words, a stronger mTOR inhibition could have a more severe impact on ER biogenesis without further reduction of the SASP. Accordingly, we tested whether the production of SASP factors contributed to making senescent cells more sensitive than proliferating cells to ER size reduction by VPF or Torin1. First, we performed single-cell RNA-seq transcriptomic analysis of senescent (ETIS) WI-38 cells treated with VPF for 72 h, after the most sensitive cells from the population had been eliminated by apoptosis, to evaluate whether the highest SASP-secreting cells were also the most vulnerable (**Fig. 3L**). We also sequenced untreated ETIS WI-38, integrated both datasets, and clustered them for comparison (**Fig. 3M**). When we evaluated the expression of 132 SASP-related mRNAs (STAR Methods) for each of the defined clusters, we observed that, in most cases, after VPF selection, there was a marked reduction in both the expression levels (1.1- to 4.2-fold decrease) and the percentage of cells expressing these transcripts (8-30.7% decrease) (**Fig. 3N**). These observations suggest that, after VPF treatment for 72 h, the remaining cells displayed a much lower secretory phenotype in all the clusters. As expected, the same VPF treatment elevated the expression of ER stress response markers in most of the clusters as well, in line with our previous findings (GSE221104, token wpexgsycrbqhdcn; GSE221117, token ilgpcwmurpibfwn). Therefore, high SASP-expressing senescent cells appeared more sensitive to VPF-triggered senolysis.

Second, to validate whether the sensitivity to VPF treatment was truly dependent on the SASP, we lowered NF-κB activity by silencing RELA (Birch and Gil, 2020; Freund et al., 2011) and evaluated the effect of VPF treatment on ER stress and senolysis. In keeping with earlier findings that NF-κB repression decreased ER stress markers in senescence (Dorr et al., 2013), EIF2A phosphorylation was reduced after RELA silencing in senescent cells (**Fig. 3O**). Importantly, silencing RELA diminished EIF2A phosphorylation triggered by treatment of senescent cells with VPF or Torin1 (**Fig. 3O** and **Fig. S3I**), and also mitigated cell death and caspase 3/7 activity (**Figs. 3P, Q** and **Fig. S3J**). As expected, silencing RELA reduced the levels of mRNAs encoding prominent SASP factors like IL1A, IL6, or IL8 (**Fig. S3K**), and lowered the levels of key SASP factors in conditioned media (**Fig. S3L**). In sum, these findings suggest that senescent cells are more sensitive to VPF (and possibly other ER-targeting drugs) because they actively producte SASP factors. We propose that senescent cells cope with SASP-triggered ER stress by activating YAP-TEAD, which in turn represses DDIT4, an essential step to retain mTOR function and preserve a robust ER biogenesis. Disruption of this delicate balance by inhibiting YAP-TEAD (using VPF) leads to excessive ER stress and subsequent apoptosis (**Fig. 3R**).

### VPF treatment reduces senescent cell burden *in vivo* in mice

In light of our findings using VPF in cultured cells, we set out to evaluate whether VPF treatment might also cause senolysis in models of senescence *in vivo* such as natural aging or doxorubicin (Doxo)- induced senescence (Anerillas et al., 2022a; Baar et al., 2017; Demaria et al., 2017). To study if VPF might influence the burden of senescent cells in naturally aging tissues, mice were treated with VPF monthly, starting at 22 months old (m.o.) until they reached 24 m.o.; in parallel, mice received either vehicle (DMSO) or the known senolytic ABT-737 following published regimens (Ovadya et al., 2018). One week after the last round of treatments, kidney, liver, and lung samples were collected (**Fig. 4A**). Overall, old mice treated with three rounds of either VPF or ABT-737 looked healthier than DMSO-treated controls (**Fig. 4B**). Importantly, in old mice treated with VPF or ABT-737 we observed a substantial decrease in the presence of the senescence markers p16 and p21 by immunofluorescence analysis (**Figs. 4C,D**) and *p16* and *p21* mRNAs by RT-qPCR analysis (**Fig. 4E**) in these tissues. Given that cell senescence exacerbates fibrosis in certain organs (Mylonas et al., 2021; Ogrodnik et al., 2017; Schafer et al., 2017), we evaluated collagen deposition by staining with Masson’s trichrome (MTC) the different groups included in this study. As shown (**Figs. 4F, G**), the elevated MTC staining observed as a result of aging was diminished with either VPF or ABT-737 treatments, in accordance with reductions in the numbers of senescent cells caused by each drug. Moreover, increased serum urea, a marker of kidney dysfunction in aged mice (Anerillas et al., 2022a; Baar et al., 2017), returned to normal levels with either treatment (**Fig. 4H**), in line with the observed reduction in kidney fibrosis.

**Figure 4.**
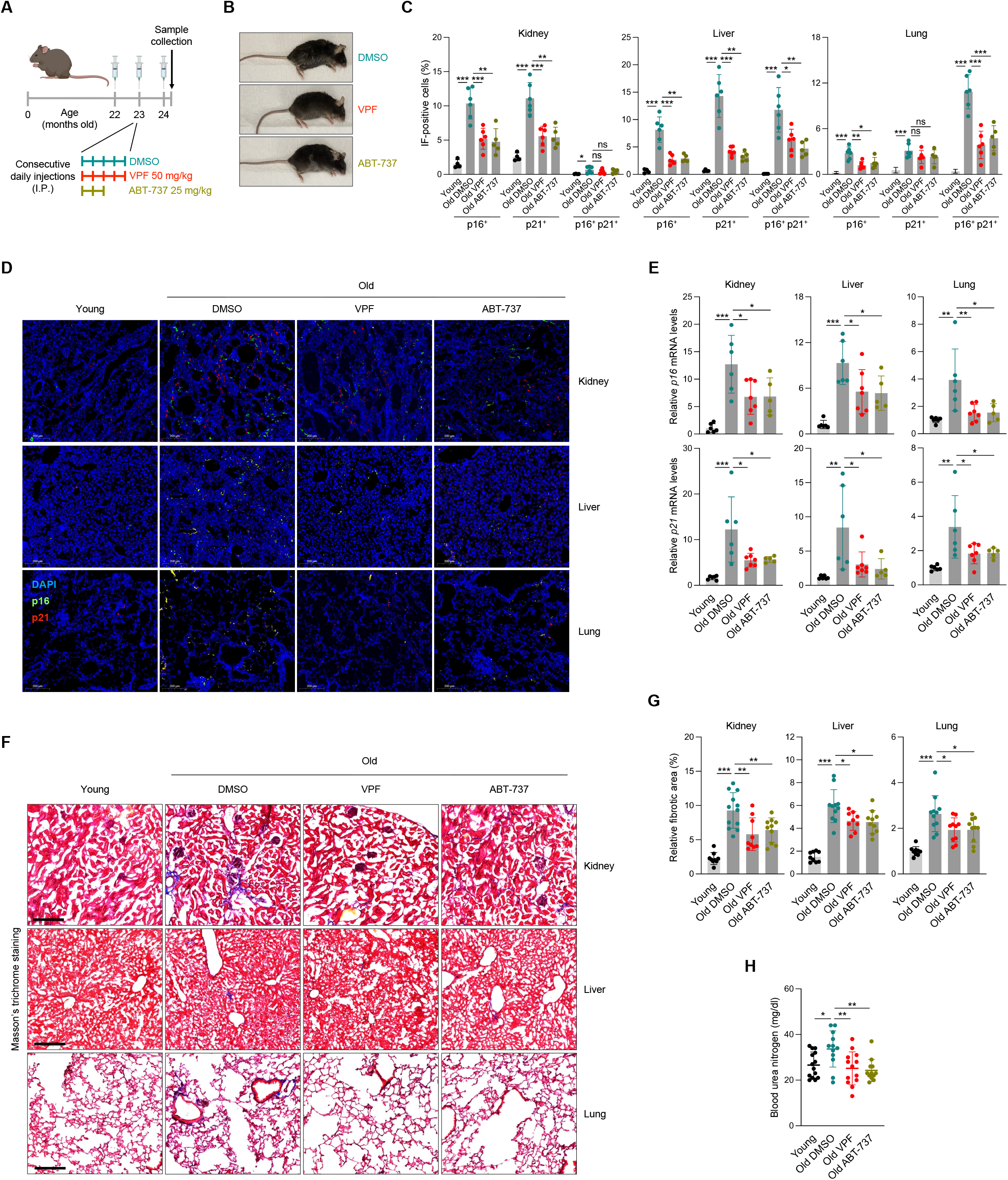
VPF treatment reduces senescent cell burden in vivo. **(A)** Schematic displaying the treatment regimens followed for each of the experimental groups included in this study (DMSO, VPF 50 mg/kg, ABT-737 25 mg/kg) in naturally aged mice. **(B)** Representative images of the general appearance of the different treatments described in (A) at the end of the study. **(C)** Quantification of the percentage of p16-, p21-, or p16/p21 double-positive cells in kidney, liver, and lung tissues for the groups described in (A). **(D)** Representative images of the data displayed in (C). **(E)** RT-qPCR analysis of *p16* and *p21* mRNA levels (normalized to *Actb* mRNA) in kidney, liver, and lung in the conditions described in (A). **(F, G)** Masson’s trichrome (MTC) staining performed in the tissues and conditions analyzed in (C), representative images of MTC staining (F) in blue, and quantification of the blue area present at each sample divided by the total area in red (G). Scale bar, 200 μm. **(H)** Serum analysis of blood urea nitrogen (mg/ml) levels for the groups described in (A). Data in (C, E, G, H) represent the means and the individual values as dots ± SD of at least n=6 independent replicates; significance (*P< 0.05, **P< 0.01, ***P< 0.001) was calculated using one-way ANOVA. See also Fig. S4.

Regarding Doxo-induced senescence *in vivo*, mice were injected with Doxo (or vehicle DMSO) to trigger senescence systemically, and VPF was administered daily for the final 4 days (**Fig. S4A**). Samples were collected at day 14 after the start of the Doxo treatment. The levels of senescence were evaluated in the lung, kidney, and liver of these mice by quantifying both *p21* mRNA levels (**Figs. S4B**) and p21-positive cells (**Figs. S4C, D**) in these organs, along with GDF15 levels in serum (**Fig. S4E**). VPF treatment markedly reduced the levels of all the senescence markers evaluated in this system, in agreement with the observations in aged mice. In conclusion, the data presented here support the notion that VPF treatment is a promising strategy to reduce the burden of senescent cells in detrimental scenarios such as aging or chemotherapy-induced senescence.

## DISCUSSION

This study uncovers a vulnerability of senescent cells linked to a unique metabolic feature, namely the robust secretion of SASP factors. As suggested previously (Dorr et al., 2013), the ER of senescent cells is subject to stress due to high demands of protein synthesis required to implement the SASP (Chen et al., 2015; Hernandez-Segura et al., 2018). Our results show that reducing ER biosynthesis by inhibiting production of PtdCho causes a metabolic crisis that primes senescent cells to die by apoptosis. Although VPF treatment was previously found to cause ER stress (Wang et al., 2020), here we have connected the derepression of DDIT4 by YAP-TEAD inhibition to the reduced mTOR function and subsequent impairment of ER biogenesis.

Senescent cells are believed to reprogram their metabolism to cope with their heightened secretory needs (Salama et al., 2014). Organelles such as the ER, the Golgi, and lysosomes appear to require efficient coordination to support the SASP (Narita et al., 2011). Current evidence suggests that mTOR is needed to support the SASP early in senescence (Herranz et al., 2015; Laberge et al., 2015); therefore, mTOR inhibition was proposed as a strategy to reduce SASP factor release by senescent cells (Soto-Gamez and Demaria, 2017). However, we found instead that higher doses of mTOR inhibitors led to senolysis, as reported (Carroll et al., 2017), and further mTOR inhibition did not lower SASP factor secretion, but instead decreased ER biogenesis by lowering the levels of Lipin1 and CCTα, two key enzymes in PtdCho biogenesis (Brandt et al., 2018; Peterson et al., 2011; Quinn et al., 2017).

The apoptotic cell death triggered by the VPF inhibition of YAP-TEAD in senescent cells suggests that YAP-TEAD functions as a pro-survival pathway in senescence, recapitulating the pro-survival effects of this pathway in other cell responses (Moya and Halder, 2019). We previously found that neurofibromin 2 (NF2), an upstream activator of the Hippo signaling cascade that suppresses YAP-TEAD, is one of the core proteins differentially phosphorylated when comparing cells committed to senescence versus apoptosis in response to etoposide (Anerillas et al., 2022b). However, an EMT-like phenotype crucial for implementing senescence over apoptosis (Anerillas et al., 2022b) seems to be suppressed by VPF treatment (**Fig. S2C**), as we and others have found (Chan et al., 2022). Thus, VPF might promote apoptosis not only by triggering ER stress but also by suppressing an EMT program.

Finally, we propose that, along with the recent discovery that senescent cells suppress the pro-apoptotic activity of p53 to remain viable (Baar et al., 2017; Sturmlechner et al., 2022), YAP-TEAD counteracts the pro-apoptotic side of p53 in senescent cells by transcriptionally repressing DDIT4 production. Therefore, forced p53 increase, for instance by using known MDM2 inhibitors, could offer better outcomes combined with YAP-TEAD inhibition when intending to achieve senolysis (van Deursen, 2019). In sum, we have identified YAP-TEAD as a signaling complex that allows senescent cells to remain viable by lowering ER stress through DDIT4 repression. By inhibiting this pathway, senescent cells can be selectively eliminated when their removal is advantageous.

## ACKNOWLEDGEMENTS

This research was supported in its entirety by the NIA IRP, NIH.

## STAR METHODS

### Cell culture and treatment

Human fibroblasts IMR-90 (ATCC), WI-38 (Coriell Institute), and BJ (ATCC) were cultured in Dulbecco’s modified Eagle’s medium (DMEM, Gibco) supplemented with 10% heat-inactivated fetal bovine serum (FBS), 0.5% Penn/Strep, Sodium Pyruvate, and non-essential amino acids (all from Gibco) in a 5% CO2 incubator. HUVEC (human umbilical vein endothelial cells) and HSAEC (primary human lung small-airway epithelial cells) were cultured in their respective media (Vascular Cell Basal Medium plus Endothelial Cell Growth Kit-BBE, Airway Epithelial Cell Basal Medium plus Bronchial Epithelial Cell Growth Kit, ATCC), supplemented with 0.5 % Penn/Strep, Sodium Pyruvate, and non-essential amino acids; and cultured under the same conditions (5% CO2 incubator). Cells were kept at low population doubling levels (PDL) for all the experiments in this study, unless indicated.

Cellular senescence was triggered as follows. Etoposide-induced senescent (ETIS) WI-38, BJ, and IMR-90 cells were cultured for 8-10 days in the presence of the different etoposide concentrations (50, 25, and 50 μM, respectively) (Etoposide, Selleckchem). For ionizing radiation-induced senescence (IRIS), WI-38 cells were exposed to 15 Gy, medium was refreshed, and cells were cultured for an additional 8-10 days. Replicative senescence (RS) was achieved after repeated passaging of WI-38 cells until they reached replicative exhaustion (PDL ~55). For HUVECs and HSAECs ETIS, cells were treated with etoposide (at 10 and 20 μM, respectively) for 3 days and media was refreshed (without etoposide) until day 8. All the remaining drugs and compounds used were refreshed every 48 h.

Transfection of siRNAs was carried out by following the manufacturer’s instructions (RNAiMAX, Invitrogen). In short, cells at 50% confluency were transfected with ON-TARGETplus SMARTPool (Dharmacon) non-targeting siCtrl (Catalog ID: D-001810-10-05), siMOB1A (Catalog ID: L-021097-00-0005), siTEAD2 (Catalog ID: L-012611-01-0005), siYAP1(Catalog ID: L-012200-00-0005), siMAP4K1 (Catalog ID: L-003586-00-0005), siTAOK2 (Catalog ID: L-004171-00-0005), siPERK (Catalog ID:L-004883-00-0005), siDDIT4 (Catalog ID: L-010855-01-0005), siTP53 (Catalog ID: L-003329-00-0005), and siRELA (Catalog ID: L-003533-00-0005) siRNAs at a final concentration of 25 nM. Twenty-four hours later, additional treatments were initiated as indicated. Cell viability was measured by direct cell counting and represented as the percentage of remaining cells compared to the number of cells present at the beginning of the experiment. Cell counts were performed manually by using ImageJ and performed in at least three independent replicates. Three fields were randomly selected and counted for each replicate analyzed.

### CRISPR screen analysis

Etoposide-induced senescent WI-38 fibroblasts (10 days in 50 μM etoposide) were transduced with the whole-genome lentiviral CRISPR/Cas9 knockout Brunello library (Addgene) at a previously-optimized MOI of 0.44 (for 16 h with 1 μg/ml polybrene). Three days later, a replicate was collected and considered as t=0. After incubation for an additional 14 days, the t=14 samples were collected, in triplicate. A total of 40,500,000 cells transduced with 18,000,000 lentiviral particles were collected for each of the replicates. These samples were sequenced on a NextSeq 500 instrument and aligned to the sgRNA sequences contained in the Brunello CRISPR library (NGS sequencing, Cellecta). sgRNA read counts were analyzed with DESeq2 package v1.32.0 (Love et al., 2014) to calculate differential sgRNA representation and statistical significance. The sgRNAs decreasing at least 2-fold (p-value < 0.05) in t=14 compared to t=0 were included in the analysis through the Enrichr platform (Kuleshov et al., 2016).

### BrdU incorporation

Cells were incubated with (4 μg/ml) 5-Bromo-2’-deoxyuridine (BrdU) diluted in the appropriate media 24 h. BrdU incorporation was assayed using a commercial kit (BrdU Cell Proliferation Assay Kit, Cell Signaling Technology, 6813) following the manufacturer’s protocol.

### Caspase 3/7 activity

Caspase 3/7 activity was measured by using the Caspase-Glo 3/7 Assay System (Promega). Cells were lysed in Caspase-Glo® 3/7 solution directly on the plate, shaken vigorously for 30 sec and incubated at 25 °C in the dark for 30 to 180 min. Luminescence was then measured using a GloMax plate reader (Promega) and normalized to cell counts.

### RT-qPCR analysis

Tissue samples (flash frozen) or cells were lysed in either Tri-Reagent (Invitrogen) or RLT buffer (Qiagen). Tissue samples needed an extra step for their disruption with a tissue homogenizer (Bertin Instruments). The resulting lysates from either tissues or cells were processed with the QIAcube (Qiagen) to purify total RNA, and then reverse-transcribed (RT) to synthesize cDNA using Maxima reverse transcriptase (ThermoFisher Scientific) and random hexamers. Real-time, quantitative (q)PCR analysis was then performed using SYBR Green mix (Kapa Biosystems), and the relative mRNA levels were determined by the 2^-ΔΔCt^ method. All mRNAs evaluated were normalized to human *ACTB* mRNA or mouse *Actb* mRNA levels. The primers used for specific detection of human mRNAs, forward (F) and reverse (R), were the following:

GTTACGGTCGGAGGCCG and GTGAGAGTGGCGGGGTC for *p16/CDKN2A* mRNA; AGTCAGTTCCTTGTGGAGCC and CATGGGTTCTGACGGACAT for *p21/CDKN1A* mRNA; AGTGAGGAACAAGCCAGAGC and GTCAGGGGTGGTTATTGCAT for *IL6* mRNA; CATGTACGTTGCTATCCAGGC and CTCCTTAATGTCACGCACGAT for *ACTB* mRNA; GACCTCAACGCACAGTACGAG and AGGAGTCCCATGATGAGATTGT for *PUMA* mRNA; CCTCAGCATCTTATCCGAGTGG and TGGATGGTGGTACAGTCAGAGC for *TP53* mRNA; ATGTGGAGATCATTGAGCAGC and CCTGGTCCTGTGTAGCCATT for *RELA* mRNA; ACCAAGCCGGATTTGCGATT and ACTTGCACTTGTTCCTCGTGG for *PMAIP1* mRNA; GCCCCACAACAAAAGAGGTC and AGGTCATTCCAGTGAGTGCTA for *TNFRSF10B* mRNA; TAGCCCTGCGTAGCCAGTTA and TCATGCTTAGTCCACTGTCTGT for *YAP1* mRNA; GCCTCCGAGAGCTATATGATCG and TCACTCCGTAGAAGCCACCA for *TEAD2* mRNA; GTCGTGGACCCTGACATTTTC and CCTTAAAGACTTCCCCATACGTG for *MAP4K1* mRNA; CAGCAGCCGCTCTTCTAAAAC and CCTCAGGCAACATAACAGCTTG for *MOB1A* mRNA; GGGAAGCAGTCCAATGAGAAA and CCGGTACTGAATGGTGTTGGG for *TAOK2* mRNA; CTCCATTGCTGAGACGTCAA and ATAGACATGCCGCCCTTCTT for *TGFB2* mRNA; AGTAGAGGAACTGGTCACTGG and TGTTTCTCGCTTTTCCACTGTT for *ANKRD1* mRNA; TGAGGATGAACACTTGTGTGC and CCAACTGGCTAGGCATCAGC for *DDIT4* mRNA; GGAAACAGAGTGGTCATTCCC and CTGCTTGAGCCGTTCATTCTC for *DDIT3* mRNA; GGAAGACAGCCCCGATTTACT and AGCACGAACTGTTGTAATGTCA for *ASNS* mRNA; GCCTTGCTTACCTTATGAGGAC and GGGAGAGATGATGCTTCGCC for *DUSP1* mRNA; TGCCGCACTCAGTGTTGTTAG and GCAATTCCCGCACAAGATTCT for *PSAT1* mRNA; GATCAGCCCCATGTTCAAAGG and GTTGGAGGCGTCCAGTACAC for *SLC6A9* mRNA; GCCATCATGCCGTAGCATC and AGCCTCAGTTCCATCACAGAT for *PCK2* mRNA; CTGCTACGGCCTTATCAGCTT and CGCTCCACATCTGCACGAA for *OXTR* mRNA; ACCCTCGCATTGACCTTCC and TTTGTGCAAAATCCCGTTGTTT for *COL11A1* mRNA; GGAAACGAGAGCCGGATTTATT and ACTATGTCCATTATGGCAGCTTC for *PERK* mRNA; ACTGCCCAAGATGAAGACCA and CCGTGAGTTTCCCAGAAGAA for *IL1A* mRNA; TCCTGATTTCTGCAGCTCTGT and AAATTTGGGGTGGAAAGGTT for *IL8* mRNA; AGCTTGCCTCAATCCTGCATCC and TCCTTCAGGAACAGCCACCAGT for *CXCL1* mRNA; GGCAGAAAGCTTGTCTCAACCC and CTCCTTCAGGAACAGCCACCAA for *CXCL2* mRNA; CAGCCAGATGCAATCAATGCC and TGGAATCCTGAACCCACTTCT for *CCL2* mRNA; TCCTGAACCTGAGTAGAGACAC and TGCTGCTTGTAGTGGCTGG for *CSF2* mRNA; TGGCGAGCAGGAGTATCAC and AGGTCTCCATCTGACTGTCAAT for *CSF1* mRNA; AGGGCAGAATCATCACGAAGT and AGGGTCTCGATTGGATGGCA for *VEGFA* mRNA; CCAACGTGACGGACTTCCC and TACACGACTATGCGGTACAGC for *LIF* mRNA; CATCACCCAGGTCAGCAAG and GCTCAAGTACACCTGGGCAC for *IGFBP7* mRNA; GGCTTGACATCATTGGCTGAC and CATTGGGCCGAACTTTCTGGT for *BDNF* mRNA; ACTACCAGAAACGAGTGGGAA and GCATCTGTTCTCGGAAAACCT for *BMP2* mRNA; ATGATTCCTGGTAACCGAATGC and CCCCGTCTCAGGTATCAAACT for *BMP4* mRNA; CAGCATGGACGTTCGTCTG and AACCACGGTTTGGTCCTTGG for *CTGF* mRNA; CTCGATCCGCTCCTTTGATGA and CGTTGGTGCGGTCTATGAG for *PDGFB* mRNA; GGAGAAGAGCGACCCTCAC and AGCCAGGTAACGGTTAGCAC for *FGF2* mRNA; TCCTGCCAACTTTGCTCTACA and CAGGGCTGGAACAGTTCACAT for *FGF7* mRNA; GACCCTCAGAGTTGCACTCC and GCCTGGTTAGCAGGTCCTC for *GDF15* mRNA; CTTCCAGCCGAGGTCCTT and CCCTGGACACCAACTATTGC for *TGFB1* mRNA; and GCCAGTATCAATTCCGACATCG and TCACCGCGTATGTGAAGGC for *WNT5A* mRNA.

The primers used for mouse transcripts, each forward (F) and reverse (R) were: CCCAACGCCCCGAACT and GCAGAAGAGCTGCTACGTGAA for *p16/Cdkn2a* mRNA; TTCTTTGCAGCTCCTTCGTT and ATGGAGGGGAATACAGCCC for *Actb* mRNA; and TTGCCAGCAGAATAAAAGGTG and TTTGCTCCTGTGCGGAAC for *p21/Cdkn1a* mRNA.

### YAP-TEAD luciferase assay

Cells were transduced at an MOI of 5 with a commercial lentiviral construct containing a luciferase reporter for TEAD activity (TEAD Luciferase Reporter Lentivirus, BPS Bioscience). Luciferase activities were measured by using a commercial kit (Steady-Glo® Luciferase Assay System, Promega). Activity measurements were normalized to cell counts.

### Co-immunoprecipitation

To perform co-immunoprecipitation analysis of the interaction between YAP and TEAD, we lysed 3 million cells per replicate with RIPA buffer plus Protease/Phosphatase Inhibitor Cocktail (Cell Signaling Technology) on ice. Samples were then sonicated twice for 20 sec, centrifuged for 10 min at 10,000 × g at 4°C, and the resulting supernatant was transferred into a new tube. One milligram of protein from each sample was incubated for 16 h at 4°C (in rotation) with either TEAD antibody (Cell Signaling Technology, 13295) or an IgG isotype control (Cell Signaling Technology, 3900), using 0.5 μg of each antibody along with 250 μg of Pierce™ Protein A/G Magnetic Beads (ThermoFisher) per condition. After 3 rounds of washes, samples were eluted in SDS lysis buffer and run as a regular Western blot. The antibodies used for immunoblotting were YAP (Cell Signaling Technology, 14074) and TEAD (Cell Signaling Technology, 13295).

### Western blot analysis

Protein extracts were obtained by lysing the samples with 2% sodium dodecyl sulfate (SDS) (Sigma-Aldrich) in 50 mM HEPES. Lysates were boiled for 5 min and sonicated, and whole-cell protein extracts were size-separated by electrophoresis using polyacrylamide gels and transferred to nitrocellulose membranes (Bio-Rad). The membranes were then blocked for 1 h with 5% non-fat dry milk and immunoblotted with the different antibodies included in this study. The primary antibodies recognized phosphorylated AKT (Ser473) (Cell Signaling Technology, Ref. 4060S), ACTB (β-Actin C4, Santa Cruz Biotechnology, sc-47778), phosphorylated EIF2A (S51) (Abcam, ab32157), ATF4 (Cell Signaling Technology, 97038), phosphorylated p70 S6K (Thr389) (Cell Signaling Technology, 9234), phosphorylated mTOR (Ser2448) (Cell Signaling Technology, 5536), XBP1s (Cell Signaling Technology, 12782), ATF6 (Cell Signaling Technology, 65880), phosphorylated YAP (S127) (Cell Signaling Technology, 13008), phosphorylated MOB1 (Thr35) (Cell Signaling Technology, 8699), MOB1 (Cell Signaling Technology, 13730), Lipin1/LPIN1 (Cell Signaling Technology, 5195), and CCTα/PCYT1A (Cell Signaling Technology, 6931). After incubation with the appropriate HRP-conjugated secondary antibodies (Jackson Immunoresearch), the chemiluminescent signals were obtained by using the Chemidoc system (Bio-Rad).

### ER tracker staining

ER-Tracker™ Red (BODIPY® TR glibenclamide) was purchased from ThermoFisher. Briefly, cells were incubated with 0.5 μM ER-Tracker™ Red for 30 min and fixed with PFA 4%; pictures were taken with a fluorescence microscope (BZ-X Analyzer, Keyence). The relative ER-positive areas were measured with ImageJ.

### Mice

All mouse work, including the import, housing, experimental procedures, and euthanasia, was approved by the Animal Care and Use Committee (ACUC) of National Institute on Aging (NIA). C57BL/6J mice were provided standard chow ad libitum and maintained under a 12:12 h light:dark cycle. All the drugs included in the study were delivered intraperitoneally, using a maximum volume of 100 μl. Doxorubicin-induced senescence was triggered after a single dose of doxorubicin (10 mg/kg), and studied>10 days later. DMSO, VPF (50 mg/kg), or ABT-737 (25 mg/kg) were used in mice treated with doxorubicin or 22-month-old mice following different regimens as specified (Results).

### Mice tissue imaging

For tissue histology, the samples were immediately fixed in 4% PFA in PBS at 4°C for 16 h. The next day, samples were cryoprotected in a 30% sucrose solution in PBS at 4°C for 16 h, embedded in OCT compound, and stored at −80°C. Samples were then cut and mounted, and the different downstream assays were performed. Immunofluorescence staining was performed with antibodies that recognized phosphorylated EIF2A (S51) (Abcam, ab131505), CDKN2A/p16INK4a (Abcam, ab54210), or p21 (Abcam, ab107099). Collagen deposition was evaluated by staining the specified tissues with Masson’s trichrome. For fibrosis evaluation, the relative area of fibrosis was calculated by measuring the area positive for Masson’s trichrome (blue) divided by the area of total tissue (dark red). All the measurements were performed with ImageJ in a randomly selected field for each of the mice included in the analysis. All of the tissue staining procedures were performed commercially by iHisto (iHisto.io).

### Bioplex and biochemical analysis of mouse serum and conditioned medium

Mouse blood was extracted, allowed to clot for 2 h at 25 °C, and centrifuged for 20 min at 2000 × g. The serum (supernatant) was collected and frozen at −80 °C. Before the assays, serum was thawed and centrifuged at 16,000 × g for 4 min. Blood urea nitrogen was measured in a Dimension EXL Integrated Chemistry system (Siemens). A custom murine Luminex Assay kit was designed by R&D Biosystems to include GDF15 for mouse serum analysis. Conditioned medium was collected from human cell cultures 48 h after adding fresh medium and was analyzed with a custom human Luminex assay platform designed to include IL1A, IL6, CXCL1, CXCL2, CCL2, LIF, IGFBP-rp1, BDNF, BMP4, FGF basic, and GDF15. Serum or medium were diluted 1:2 using Calibrator Diluent RD6-52. Standards, blanks, and serum were incubated with the microparticle cocktail for 2 h at 25 °C, and then incubated with a biotin-antibody cocktail for 1 h. A final incubation of 30 min with Streptavidin-PE was carried out with shaking at 25 °C prior to plate analysis on the Bio-Rad Bioplex-200 instrument. Every incubation step was followed by three washes with wash buffer. The results were analyzed with the Bio-Plex Manager software.

### Detection of SA-β-Gal

Senescence-associated β-Galactosidase (SA-β-Gal) activity was assayed using a commercial kit (Senescence β-Galactosidase Staining Kit, Cell Signaling Technology, 9860) following the manufacturer’s instructions.

### Transmission electron microscopy

Cells in culture were processed for transmission electron microscopy visualization. Fixation for electron microscopy was performed at RT using 2.5% glutaraldehyde in 0.1 M sodium cacodylate buffer, pH 7. Cells were then scraped and gently spun down in 1.5-ml microcentrifuge tubes. Samples were post-fixed in 1% osmium tetroxide for 1 h at 4°C in the same buffer, dehydrated in increasing % ethanol and embedded in Embed 812 resin (Electron Microscopy Sciences, Hatfield, PA) through a sequence of resin resin-propylene oxide gradients until reaching pure resin. Blocks were formed in fresh resin contained in the same conical tubes, and the resin was polymerized for 48-72 h at 65°C. Blocks were trimmed and sectioned in an EM UC7 ultramicrotome (Leica Microsystems, Buffalo Grove, IL) to obtain both semi-thick (0.5-1 μm width) and ultrathin (40-60 nm width) sections. Semi-thick sections were mounted on glass slides and stained with 1% toluidine blue in a 1% borax aqueous solution for 2 min and visualized in a Leica AXIO Imager light microscope for quality control. Ultrathin sections were stained with uranyl acetate and lead citrate, and then imaged on a Talos L120C Transmission Electron Microscope (TEM) with a 4K Ceta CMOS camera. Maximal ER cisternae thickness per cell was measured with ImageJ by measuring the longest distance between ER cisternae walls in each cell. A disorganization score was given according to the level of disorganization shown by the ER in each cell (ranked 1 to 4). Parameters such as continuity or shape were considered to determine the score. Thirty cells were analyzed for each condition.

### ChIP-qPCR

Chromatin immunoprecipitation and subsequent qPCR were performed by using SimpleChIP® Plus Enzymatic Chromatin IP Kit (Cell Signaling) following the manufacturer’s instructions. Briefly, 10 μg of sheared chromatin (150-900 bp) were incubated overnight with 0.25 μg of either anti-YAP (Cell Signaling, 14074) or Normal Rabbit IgG (Cell Signaling, 2729). The eluted DNA was de-crosslinked, column-purified, and quantified by RT-qPCR analysis. Data were displayed by normalizing the percent of input recovered by YAP antibody to the percent of input recovered by IgG control for each replicate (not shown in the graphs). Negative control used is a primer set (ACCAACACTCTTCCCTCAGC and TTATTTTGGTTCAGGTGGTTGA) that amplifies a region of Chromosome 10 known to be unrelated to YAP-TEAD binding (Stein et al., 2015). Primers for DDIT4 gene (TGTTTAGCTCCGCCAACTCT and CACCCCAAAAGTTCAGTCGT) were designed to target a region in exon 2 that is at a distance from YAP binding sequence in the regulatory region of the *DDIT4* gene, shorter than 900 bp.

### RNA-seq

Bulk RNA was extracted with RLT buffer (Qiagen) using the QIAcube system (Qiagen) following the RNeasy plus protocol. The quality and quantity of RNA were assessed using the Agilent RNA 6000 nano kit on the Agilent Bioanalyzer. High-quality RNA (125 ng) was used for the library prep using Illumina TruSeq Stranded mRNA Library prep kit following the manufacturer’s protocol (Illumina, San Diego, CA). The quality and quantity of the libraries were checked using the Agilent DNA 1000 Screen Tape on the Agilent Tapestation. Paired-end sequencing was performed for 103 cycles with an Illumina NovaSeq sequencer. BCL files were de-multiplexed and converted to FASTQ files using bcl2fastq program (v2.20.0.422). FASTQ files were trimmed for adapter sequences using Cutadapt version v1.18 and aligned to human genome hg19 Ensembl v82 using STAR software v2.4.0j. Gene counts were generated using featureCounts (v1.6.4) software and normalized with DESeq2 package (v1.32.0). Further analysis of the transcriptomic differences among samples was performed using normalized counts in the GSEA platform (Subramanian et al., 2005). YAP-TEAD targets were identified by both existing evidence on the literature and analysis with MAGIC database (Roopra, 2020).

Single-cell RNA-seq libraries were prepared as indicated by the Chromium Next GEM Single Cell 3’ Reagent Kits v3.1 (10x Genomics) user guide. Briefly, ETIS cells (untreated and VPF-treated for 72 h) were trypsinized, washed with PBS, and resuspended in 10% FBS and 0.1 mM EDTA in PBS at a concentration of 900–1,000 cells/μl. The resulting single-cell suspension was loaded into a Chromium Next GEM Chip G (10x Genomics), and GEMs were generated with the Chromium Single Cell Controller (10x Genomics). Libraries were prepared using 11 and 13 cycles for cDNA amplification and Sample Index PCR (Single Index Kit T Set A, 10x Genomics), respectively, and were subjected to paired-end sequencing on a SP100 flow cell on an Illumina NovaSeq platform. The raw single-cell RNA-seq data were processed through Cell Ranger software 6.0.0 (10x Genomics) and sequencing reads were mapped to a pre-built human reference (GRCh38) (version 2020-A, 10x Genomics). Filtered matrix files generated by Cell Ranger were processed using R (version 4.2.2) and the Seurat R package (version 4.2.0). Cells expressing more than 200 transcripts and less than 15% of mitochondrial genes were included in the analysis. The untreated and VPF-treated samples were processed separately; both were log-normalized, scaled to 10,000, dimensionally reduced by principal component analysis (PCA) using the top 2,000 highly variable features and visualized by performing uniform manifold approximation and projection (UMAP). Both samples were then integrated using the Seurat CCA method. The integrated data were scaled, and dimensionally reduced using PCA and UMAP as above. Single cells in the integrated data were clustered into six subgroups using r=0.2 resolution. The gene set used for analyzing ER stress and was downloaded from MsigDB (GOBP_RESPONSE_TO_ENDOPLASMIC_RETICULUM_STRESS). A list of total of 132 SASP-related genes included for analysis was obtained after filtering out the secretory genes not detected by bulk RNA-seq in WI-38 cells, and were used as a broader gene set for SASP (**Supplementary Table 3**). The average expression levels of each gene set on single cells were calculated using the AddModuleScore function in Seurat. The returned module scores in single cells in each cluster were averaged and scaled, and were subsequently used for gene set activity comparison in dot plots.

Sequencing data are deposited in GSE221104 (token wpexgsycrbqhdcn) and in GSE221117 (token ilgpcwmurpibfwn).

### Statistical analysis

Data are represented as the means ± standard deviation (S.D.) of at least n=3 independent experiments (except for the ChIP qPCR experiments, done in duplicate). Individual data points are shown in all the bar plots. Significance was determined using two-tailed Student’s *t*-test (*P< 0.05, **P< 0.01, ***P<0.001) for cell culture experiments, while one-way ANOVA (*P< 0.05, **P< 0.01, ***P<0.001) was used for *in vivo* experiments. All the statistical analyses were carried out with Prism 9.

