## Supplemental Text and Figures for "The YAP-TEAD complex promotes senescent cell survival by lowering endoplasmic reticulum stress"

Running title: YAP and TEAD lower ER stress, protect senescent cells

### LEGENDS FOR SUPPLEMENTAL FIGURES

**Figure S1. Extended data on Figure 1.**

**(A)** Cell viability assessment by direct cell counting of senescent WI-38 cells treated with puromycin (1 µg/ml, 48 h) 72 h after being transduced with the Brunello library at the indicated MOIs. The gray bars represent the expected viability if the transduction efficiency was complete while the teal bars represent the viability observed for each of the MOIs after puromycin treatment.

**(B)** Cell viability as assessed by direct cell counting of WI-38 cells transfected with the indicated siRNAs and rendered senescent after treatment with etoposide for 8 days (ETIS).

**(C)** Analysis of the levels of the indicated mRNAs in proliferating (P) or ETIS WI-38 cells transfected with the indicated siRNAs 24 h before either treatment with etoposide (50 µM) or no treatment, and culture for an additional 8 days.

**(D)** Western blot analysis of the levels of phosphorylated YAP (S127), YAP, phosphorylated MOB1 (T35), MOB1, and ACTB levels at the indicated conditions.

**(E, F)** Analysis of BrdU incorporation (E) and SA-β-Gal staining (F) in the indicated cell types, rendered senescent by etoposide (ETIS), Ionizing Radiation (IRIS), or replicative exhaustion (RS).

**(G)** Caspase 3/7 activity measured in RS and IRIS WI-38 cells treated for 72 h with the indicated doses of Verteporfin (VPF).

**(H, I)** Cell viability as assessed by direct cell counting (H) and Caspase 3/7 activity measurement (I) for the indicated models of senescence along with proliferating controls, after either no treatment or treatment with VPF for 72 h at the indicated doses.

Graphs in (B, C, E, G-I) represent the means and each individual value as a dot ±SD of at least n=3 independent replicates; significance (*P < 0.05, **P < 0.01, ***P < 0.001) was determined using two-tailed Student’s *t*-test. Unless indicated, statistical tests were performed relative to untreated or proliferating controls.

**Figure S2. Extended data on Figure 2.**

**(A)** Heatmap displaying the differential expression of the indicated transcripts (by row Z-Score) in ETIS WI-38 fibroblasts that were left untreated or treated with VPF for 48 h. Proliferating untreated controls were included as baseline control.

**(B)** Representative Western blot analysis of the levels of YAP and TEAD proteins after immunoprecipitation experiments with the indicated antibodies (IgG or TEAD) in ETIS WI-38 fibroblasts that were either untreated or treated with VPF for 48 h. IgG bands are pointed out with arrows placed on the left side of the panel. Inputs are also included.

**(C)** *Left*, GSEA analysis of the association (enrichment score) with the gene set “Hallmark: Epithelial-Mesenchymal Transition” of ETIS WI-38 cells treated with VPF (48 h) compared to untreated senescent cells (-). *Right*, heatmap of the expression score of the indicated transcripts included in the same gene set.

**(D)** Western blot analysis of the levels of ATF6, XBP1s, and loading control ACTB for the conditions described in (C).

**(E)** GSEA plot showing the association (enrichment score) of the gene set “GOBP: PERK-mediated UPR” with the conditions described in (C).

**(F)** Western blot analysis of the levels of phosphorylated EIF2A (S51) and loading control ACTB in WI-38 cells transfected with siCtrl or siPERK, rendered senescent with etoposide (ETIS) and then either left untreated or treated with 1.5 µM VPF for 48 h.

**(G, H)** Cell viability assessment by direct cell counting (G) and RT-qPCR analysis of *PERK* mRNA levels (H) in the conditions described in (F).

**(I, J)** Maximal cisternae thickness (I) and disorganization score (J) as measured by transmission electron microscopy (TEM) in the groups described in (C). Thirty cells were analyzed for each condition.

**(K)** Relative binding to the regulatory region of the *DDIT4* gene or a negative control (Neg Ctrl) DNA in YAP ChIP samples of ETIS WI-38 cells that were untreated or treated with 1.5 µM VPF (48 h).

**(L)** RT-qPCR analysis of the levels of *DDIT4* and *p53* mRNAs in WI-38 cells transfected with the indicated siRNAs, rendered senescent with etoposide (ETIS) and either left untreated or treated with 1.5 µM VPF for 48 h. Proliferating WI-38 cells transfected with siCtrl were included as controls.

Graphs in (G, H, K, L) display the means and the individual values as dots ±SD of at least n=3 independent replicates; graphs in (I, J) show the means and the individual values as dots ±SD of n=30 different cells. Significance (*P < 0.05, **P < 0.01, ***P < 0.001) was calculated using two-tailed Student’s *t*-test.

**Figure S3. Extended data on Figure 3.**

**(A)** Cell viability as assessed by direct cell counting of proliferating (P) or ETIS WI-38 fibroblasts that were either left untreated or treated with 100 nM Torin1 for 48 h.

**(B)** Representative micrographs showing the differences in viability between ETIS WI-38 fibroblasts that were either left untreated or treated with Torin1 as in (A).

**(C)** Western blot analysis of the levels of ATF6, XBP1s, and loading control ACTB in the conditions described in (A).

**(D)** RT-qPCR analysis of the levels of *PUMA*, *PMAIP1*, and *TNFRSF10B* mRNAs in ETIS WI-38 cells that were either left untreated or treated with Torin1 as in (A). Untreated P cells were included as baseline controls.

**(E)** Dot plot representation of the values calculated for the ER-positive relative area per cell (60 cells per condition) for the conditions described in Fig. 3G.

**(F)** Micrographs showing the areas corresponding to the endoplasmic reticulum (ER) in red for the indicated treatments. All of the treatments were performed for 48 h. Phosphatidylcholine (PtdCho) was supplemented at 50 µM.

**(G)** Heatmaps displaying the differences in SASP mRNA levels represented by row Z-Score for each transcript in ETIS WI-38 cells that were either left untreated or treated with 1.5 µM VPF (left heatmap) or 100 nM Torin1 (right heatmap) for 48 h. Untreated P WI-38 cells were included as baseline controls.

**(H)** Heat map of the row Z-Score calculated for the differences in the secretion of the indicated SASP members among the groups described in (G), including proliferating (P) WI-38 cells as a control for baseline secretion.

**(I)** Western blot analysis of phosphorylated EIF2A (S51) and ACTB levels in WI-38 cells transfected with the indicated siRNAs, made senescent with etoposide for 8 days, and then either left untreated or treated with 100 nM Torin1 for 48 h.

**(J)** Cell viability measurement by direct cell counting of the conditions described in (I).

**(K,L)** RT-qPCR analysis (K) and Bioplex analysis of the conditioned media (L) to assess SASP production and secretion in WI-38 cells transfected with siCtrl or siRELA, and rendered senescent with etoposide for 8 days. Proliferating controls transfected with siCtrl siRNA were included.

Graphs in (A, D, E, J, K) represent the means and individual values (dots) of at least n=3 independent replicates. Significance (*P < 0.05, **P < 0.01, ***P < 0.001) was calculated using two-tailed Student's *t*-test.

**Figure S4. Extended data on Figure 4.**

**(A)** Schematic representation of the treatment regimen carried out to trigger doxorubicin-induced senescence *in vivo* in mice (10 mg/kg), along with 4 consecutive treatments with DMSO (Vehicle) or VPF (50 mg/kg) from day 10 onwards. Samples were collected at day 14 after doxorubicin treatment.

**(B)** RT-qPCR analysis of *p21* mRNA levels in lung, liver, and kidney from the groups described in (A). Untreated mice were included as baseline controls.

**(C, D)** Quantification (C) and representative images (D) of p21 immunofluorescence in the conditions described in (B). Scale bar (white) represents 200 µm.

**(E)** Serum measurement of GDF15 levels for the experimental groups described in (B).

Graphs in (B, C, E) display the means and the individual values as dots ± SD of n=6 independent replicates per group; significance (*P < 0.05, **P < 0.01, ***P < 0.001) was calculated using one-way ANOVA.

Anerillas et al, Figure S1

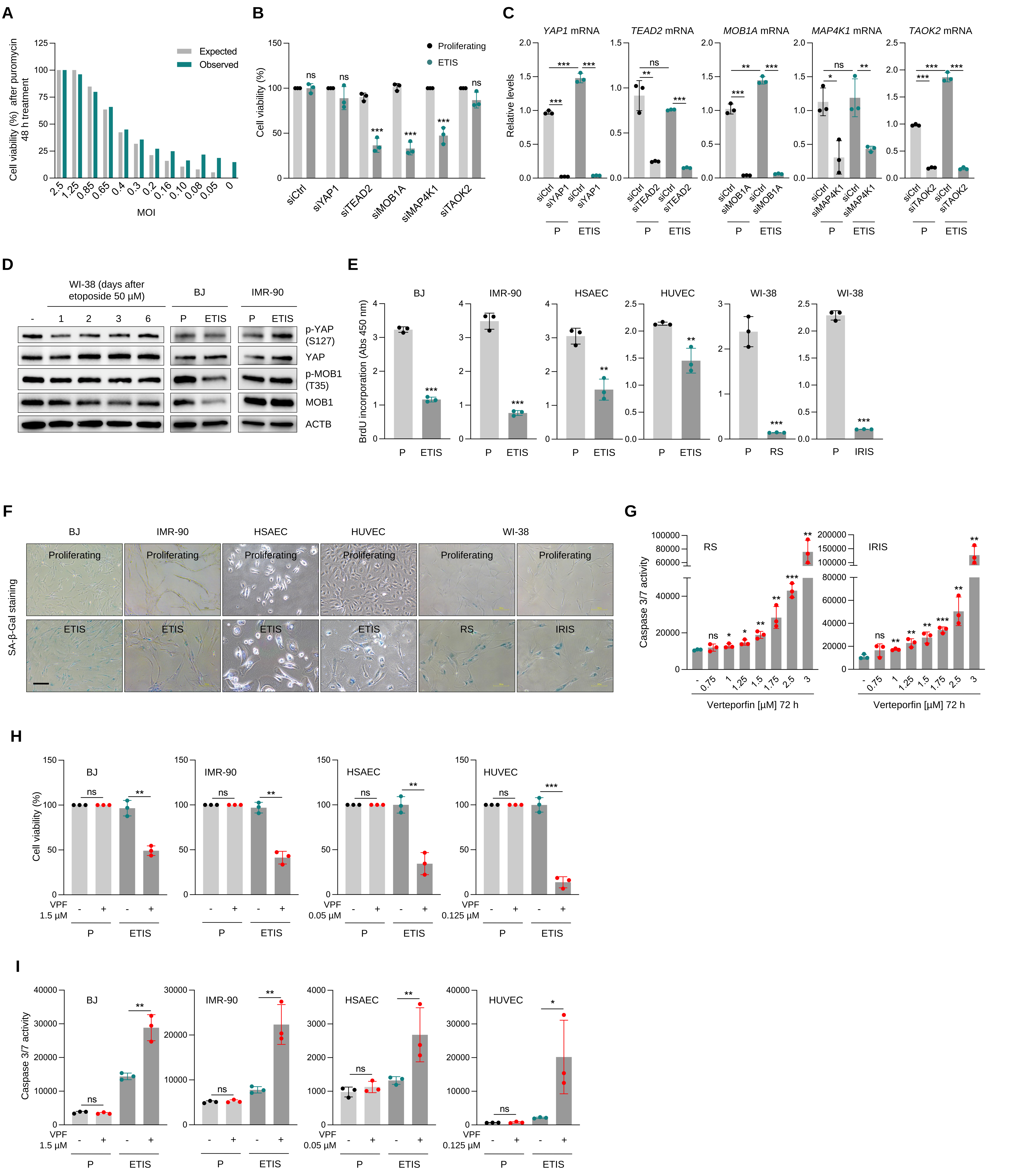

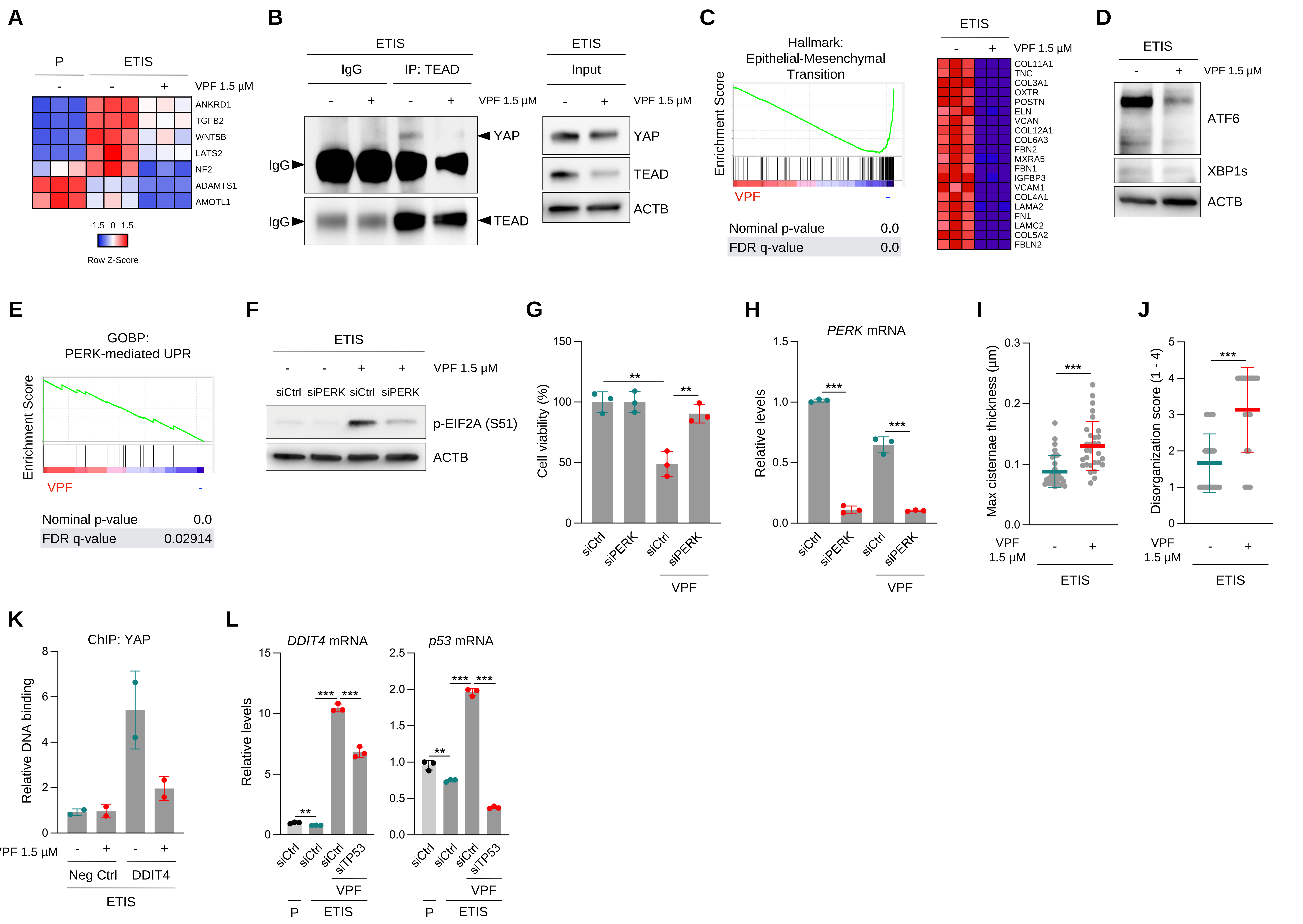
A
P
ETIS
VPF 1.5 µM
ANKRD1
TGFB2
WNT5B
LATS2
NF2
ADAMTS1
AMOTL1
Row Z-Score
B
ETIS
IgG
IP: TEAD
Input
YAP
TEAD
ACTB
C
Hallmark: Epithelial-Mesenchymal Transition
Enrichment Score
Nominal p-value 0.0
FDR q-value 0.0
D
ATF6
XBP1s
E
GOBP: PERK-mediated UPR
Nominal p-value 0.0
FDR q-value 0.02914
F
p-EIF2A (S51)
G
Cell viability (%)
H
PERK mRNA
Relative levels
I
Max cisternae thickness (µm)
J
Disorganization score (1 - 4)
K
ChIP: YAP
Relative DNA binding
Neg Ctrl
DDIT4
L
DDIT4 mRNA
p53 mRNA
siCtrl
siPERK
siTP53

Anerillas et al, Figure S3

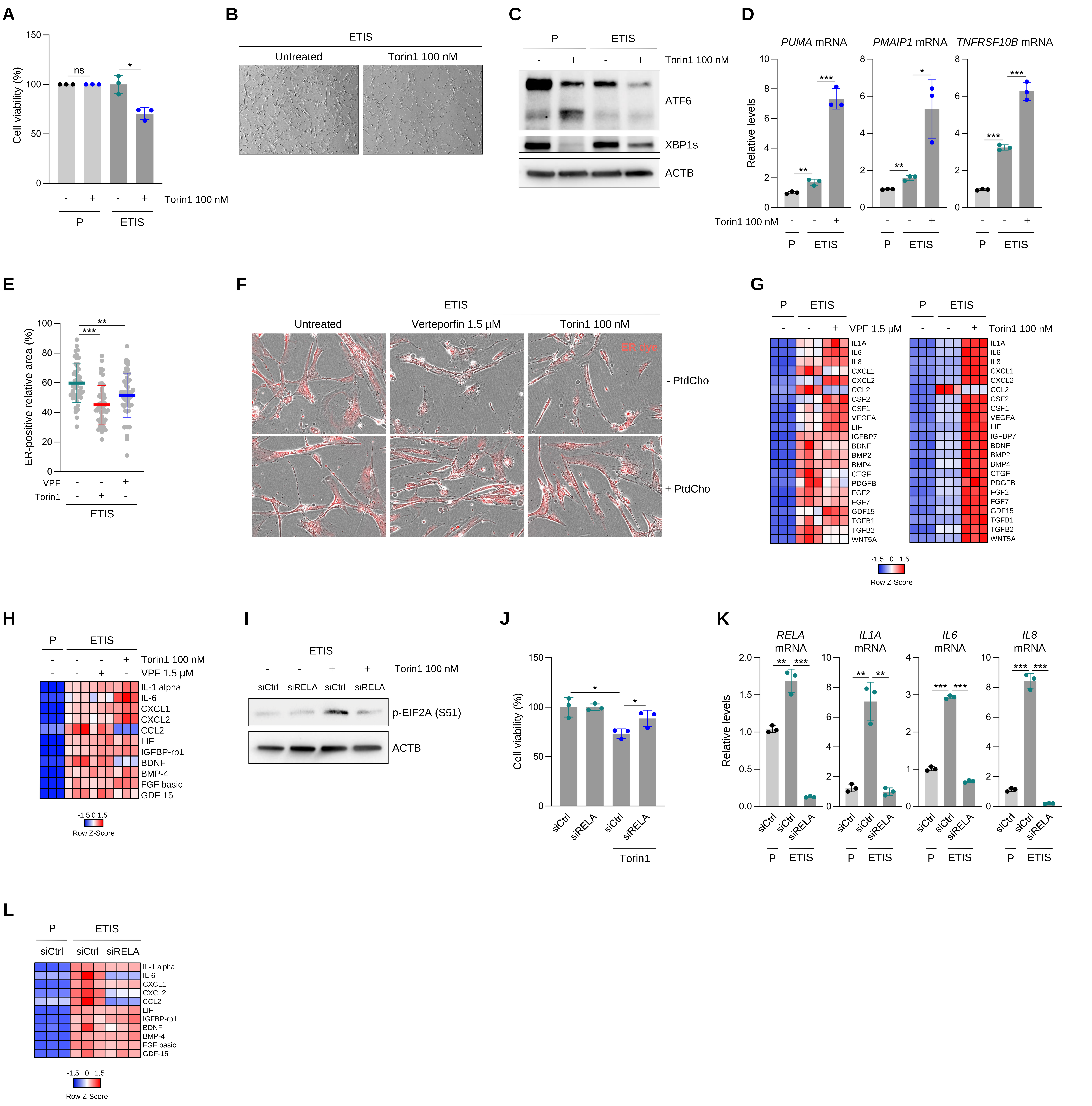

Anerillas et al, Figure S4

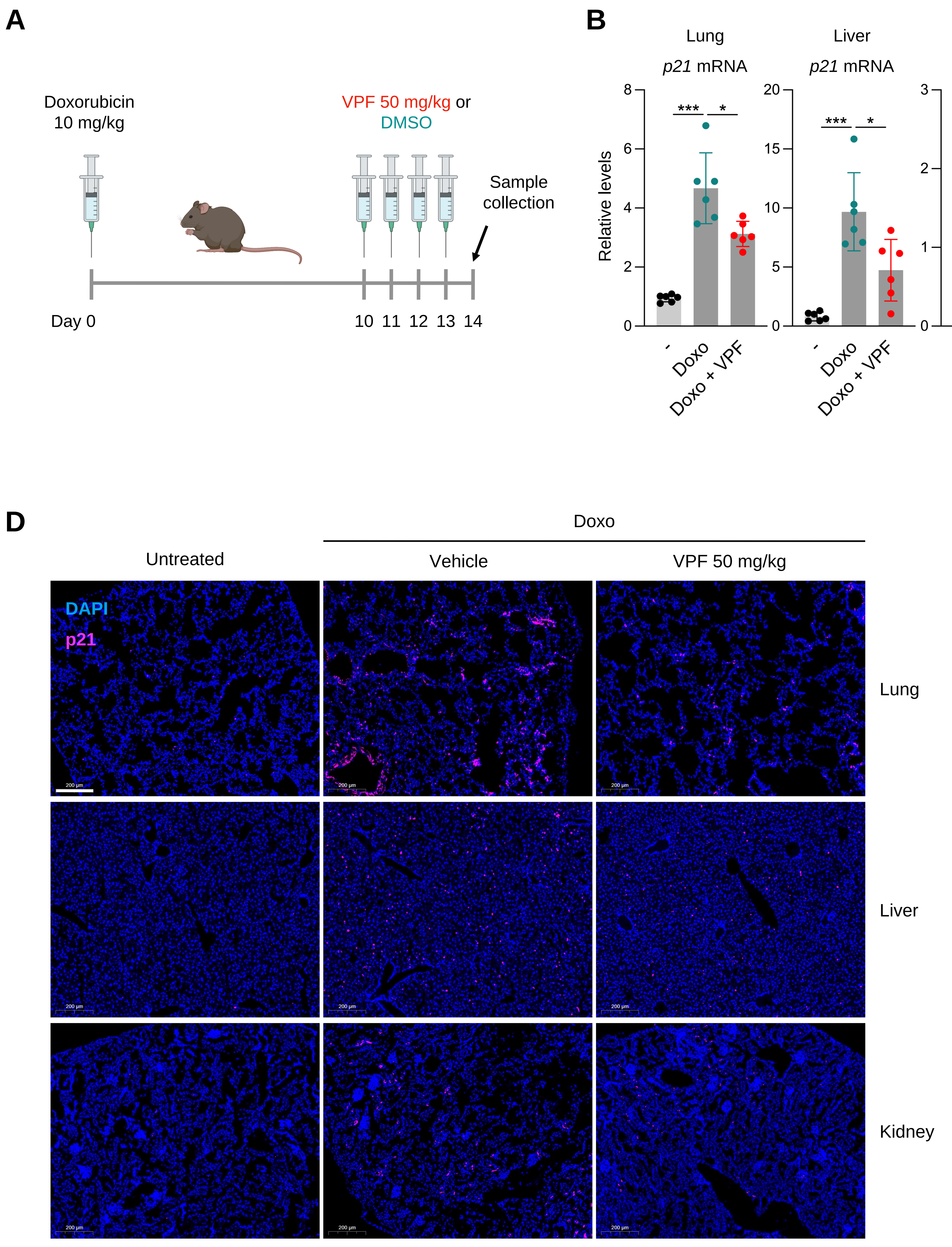
